# Genome-wide Spatial Expression Profiling in FFPE Tissues

**DOI:** 10.1101/2020.07.24.219758

**Authors:** Eva Gracia Villacampa, Ludvig Larsson, Linda Kvastad, Alma Andersson, Joseph Carlson, Joakim Lundeberg

## Abstract

Formalin-fixed paraffin embedding (FFPE) is the most widespread long-term tissue preservation approach. Here we present a procedure to perform genome-wide spatial analysis of mRNA in FFPE tissue sections. The procedure takes advantage of well-established, commercially available methods for imaging and spatial barcoding using slides spotted with barcoded oligo(dT) probes to capture the 3’ end of mRNA molecules in tissue sections. First, we conducted expression profiling and cell type mapping in coronal sections from the mouse brain to demonstrate the method’s capability to delineate anatomical regions from a molecular perspective. Second, we explored the spatial composition of transcriptomic signatures in ovarian carcinosarcoma samples using data driven analysis methods, exemplifying the method’s potential to elucidate molecular mechanisms in heterogeneous clinical samples.

## Main

Formalin-fixing and paraffin-embedding of clinical biospecimens has been the preferred method for tissue preservation for decades. FFPE is not only less expensive and easier to use than freezing-based methods, it also offers a high degree of preservation of morphological detail^1^. As a consequence, there are vast numbers of FFPE biospecimens readily available for genomics research, which could be used for extensive longitudinal studies on large patient cohorts.

However, formalin fixation negatively affects nucleic acid integrity and accessibility; due to formalin-mediated strand cleavage and formation of cross-linked adducts between RNA and other biomolecules^2^. Genome-wide quantification strategies have been developed for application on bulk FFPE samples^3,4^, which typically involve ribosomal depletion or targeted capture using oligonucleotide probe hybridization for enrichment of fragmented mRNA. Recently, several methods have been developed for spatial analysis of tissue sections (recently reviewed by Asp et al.^5^) and can broadly be characterized into 1) hybridization, which require pre-existing knowledge of the targets for probe design and 2) sequencing based approaches, which allow for unbiased mRNA poly(A) capture in the tissue being analyzed spatially. There are currently limited options for sensitive spatial transcriptomic analysis of FFPE samples^6^.

In particular, Spatial Transcriptomics^7^ has repeatedly demonstrated its value on fresh frozen (FF) tissue sections for exploration and profiling of transcriptomic landscapes^8-11^, and is today commercially available. So far, formalin-induced cross-linking and mRNA fragmentation has hampered the analysis of FFPE samples with spatially barcoded methodologies^7^.

Here, we present a protocol adapted to recover spatially-resolved mRNA from FFPE tissue sections. Recovery of mRNA for spatial analysis is achieved by removing paraffin and cross-links *in situ* with the tissue section placed on a barcoded slide (Suppl. Fig. 1). In short, tissue sections were dried, deparaffinized, hematoxylin and eosin (H&E) stained and imaged under a high-resolution microscope, followed by cross-link reversal. Decrosslinking is performed by heat-induced retrieval^12^ at 70°C with Tris and EDTA containing buffer at pH 8.0. A pH of 8.0 has previously been reported to avoid unwanted side reactions such as RNA pH-dependent hydrolysis^13^. Moreover, a quenching mechanism has been proposed between Tris and formaldehyde molecules^2^. After cross-link reversal, the tissue sections were permeabilized, reverse transcribed and sequencing libraries were generated according to the manufacturer’s protocol (see methods) with some modifications due to the short size of the mRNA molecules. Since most fragments in the final library contained the template switch oligo (TSO) sequence (Suppl. Fig. 1b), sequencing was therefore conducted with a custom primer complementary to TSO sequence, thereby reducing sequencing cost.

**Fig. 1.**
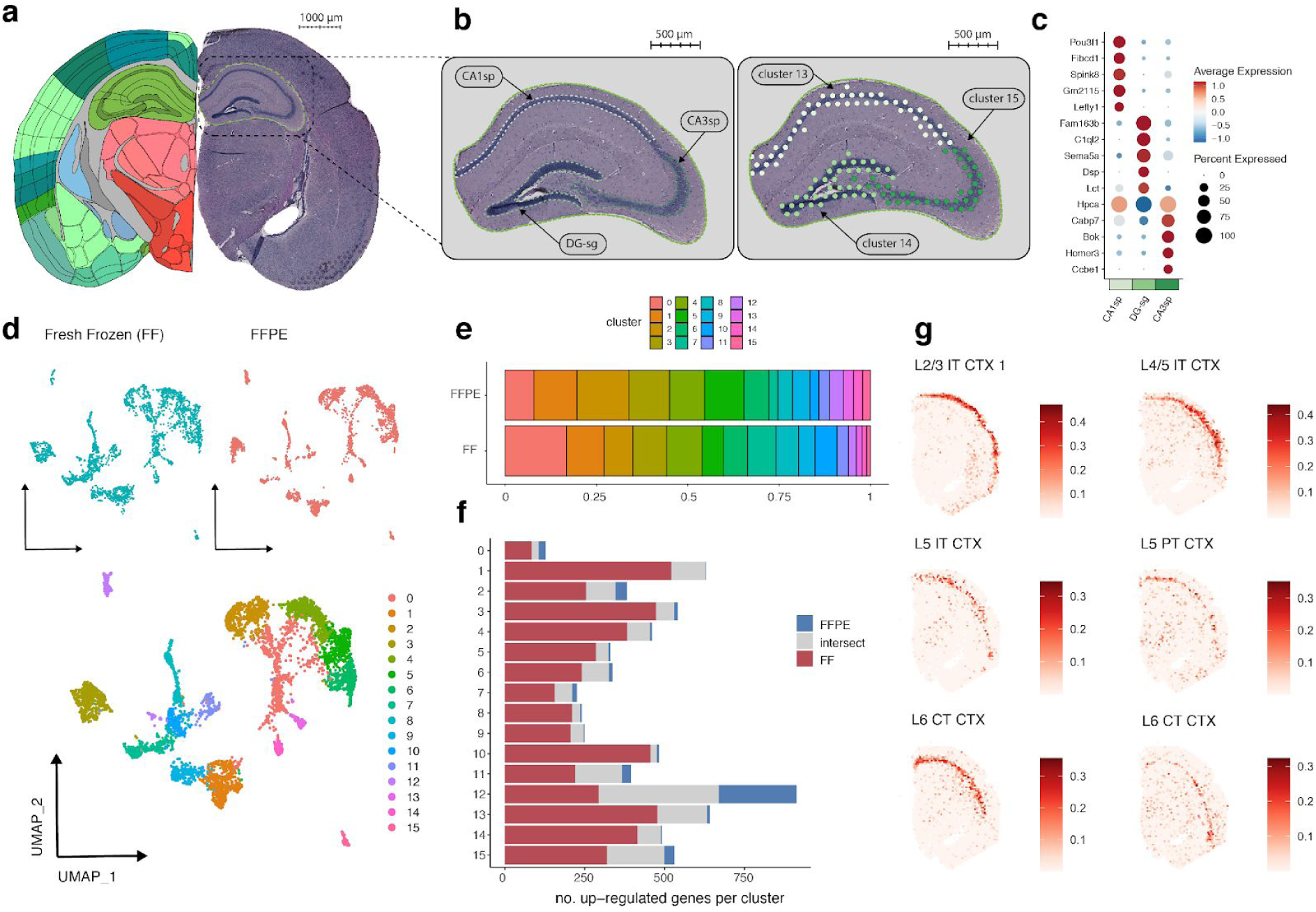
Analysis of a coronal section from FFPE mouse brain tissue. **a**, H&E image of a coronal section (right) registered to the Allen Mouse Brain Atlas (left) with the hippocampal region highlighted with a green dotted line. **b**, Zoomed in view of the hippocampal region. Three anatomical structures are highlighted (left); field 1 pyramidal layer (CA1sp), field 3 pyramidal layer (CA3sp) and the dentate gyrus (DG-sg). Three selected clusters from the unsupervised analysis (right) exemplify the correspondence between structures defined by molecular features and neuroanatomy. **c**, Dot plot visualizing the top 5 most significant marker genes for each hippocampal region in **b. d**, UMAP embedding of spots using the integrated representation (*harmony*). Spots from fresh frozen tissue (FF) colored in blue (top left), spots from FFPE tissue colored in red (top right) and spots colored by cluster (bottom). **e**, Stacked bar chart showing the relative proportions of integrated clusters in the two datasets (FFPE and FF). **f**, Stacked bar chart showing the number of marker genes detected per cluster in the FFPE and FF sections. Red bars represent genes that are exclusively found in the FF data, blue bars represent genes found exclusively in the FFPE data and grey bars represent the intersect, i.e. genes that are found to be up-regulated in both datasets. **g**, Examples of neuronal cell type subclasses from the isocortex mapped to the FFPE coronal section using *stereoscope*, with scRNA-seq (SMART-seq) data obtained from the Allen Brain Atlas. Values on the colorbar correspond to cell type proportions. Cell type subclass abbreviations; L2/3 IT CTX 1, cerebral cortex layer 2/3 intratelencephalic; L4/5 IT CTX, cerebral cortex layer 4/5 intratelencephalic; L5 IT CTX, cerebral cortex layer 5 intratelencephalic; L5 PT CTX, cerebral cortex layer 5 pyramidal tract; L6 CT CTX, cerebral cortex layer 6 corticothalamic; L6b CTX, cerebral cortex layer 6b.

The mouse brain has been extensively characterized using multiple genomics methods coupled with detailed neuroanatomy maps in projects such as the Allen Brain Atlas^14,15^. Such efforts have made the mouse brain a suitable model tissue to validate the developed approach. First, we collected a tissue section from one hemisphere of an FFPE mouse brain (coronal plane) and generated spatial expression data using our FFPE protocol. Using the *wholebrain* framework^*16*^, we registered the H&E image of the coronal tissue section to the anatomic reference of the Allen Mouse Brain Atlas (Fig. 1a). At a sequencing depth of ∼50k reads per tissue-covered spot, a total of 2533 barcoded capture locations (spots) were obtained with an average of ∼1200 unique genes and ∼2200 unique molecules detected per spot (Suppl. Table. 1). After normalization of the data, expression-based clustering revealed a total of 16 clusters (Suppl. Fig. 2a-c) which clearly corresponded to established anatomical structures.

**Fig. 2.**
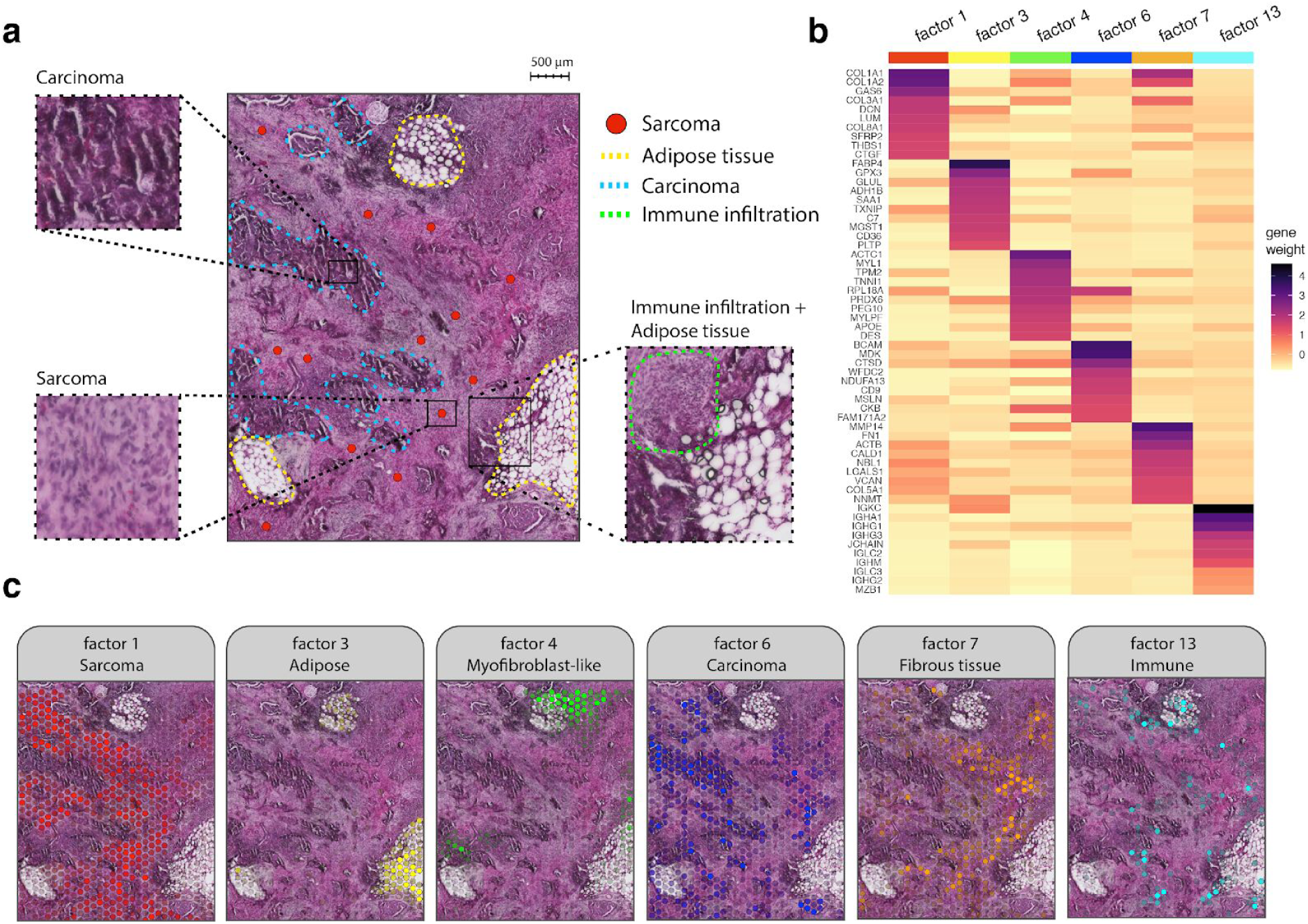
Exploratory analysis of FFPE ovarian carcinosarcoma tissue. **a**, Histological image (H&E) of a representative area manually annotated based on morphological features. Blue dotted lines delimit areas with typical slit-like morphology associated with carcinoma. Yellow dotted lines mark fat lobules dispersed across the tissue. Sarcomatous areas (red dots) are mainly located in the spaces between the fat lobules and carcinoma, displaying various degrees of malignancy. Immune infiltration (green dotted area) is also visible in certain areas. **b**, Heatmap showing the top 10 most contributing genes for a selected panel of factors (values have been rescaled within each column). **c**, Spatial activity maps of a selected panel of factors (same as in **b**), showing the link between morphological features and expression-based patterns. The factor activity values have been rescaled to range from 0 to 1. The spots with lower values are more transparent whereas spots with higher values are less transparent.

Clusters 2, 13, 14 and 15 formed the hippocampal region (HIP), whereas clusters 13 and 15 mapped to field 1 and 3 of the pyramidal layer of Ammon’s horn and cluster 14 mapped to the granule cell layer of the dentate gyrus (DG-sg) (Fig. 1b). These results exemplify how even small anatomical structures can be resolved with this approach. Moreover, differential expression (DE) analysis showed up-regulation of known marker genes *Fibcd1* and *Spink8* in the CA1sp region; *Cabp7* and *Bok* in the CA3sp; *C1ql2* and *Sema5a* in the DG-sg (Fig. 1c and Suppl. Fig. 2c). Inspection of the 2D UMAP embedding of the clustered data (Suppl. Fig. 2b) revealed larger structures in the data. These structures reflected the spatial organization of clusters forming the cerebral cortex (CTX), hypothalamus (HY), fiber tracts, thalamus (TH), striatum (STR), lateral ventricle (VL) and the hippocampal region (HIP).

For each one of the 16 clusters, one representative marker was selected from the DE analysis (Suppl. Table. 2) and its expression patterns were compared with published *in situ*^14^ images (Suppl. Fig. 3). Together, the analysis highlights that the proposed method can extract sufficient transcriptomic data from FFPE tissue sections in order to clearly and spatially identify anatomical regions of interest.

To systematically assess the quality of our FFPE-derived spatial transcriptome data, we downloaded a publicly available spatial dataset (2698 spots) from a coronal section of FF mouse brain tissue^17^, collected approximately at the same distance from Bregma along the anterior-posterior axis as the FFPE section (Suppl. Fig. 4a). At a sequencing depth ∼115k reads per tissue-covered spots, the FF dataset covered on average ∼6000 unique genes and ∼27200 unique molecules per spot, almost 5x more unique genes and 12x more unique molecules than recovered from the FFPE tissue section (Suppl. Fig 4b). Most of the captured molecules were annotated as protein coding in both datasets, but varied considerably in relative abundance of lincRNA and ribosomal protein coding transcripts (Suppl. Fig. 4c). The Pearson r-score between the FFPE and FF datasets treated as bulk was 0.95 (Suppl. Fig. 4d), indicating a strong correlation between the datasets despite a substantial difference in gene recovery.

Next, we applied *harmony*^18^ to integrate the two datasets and re-clustered the data to find a total of 16 integrated clusters shared across the two conditions (Fig. 1d-e and Suppl. Fig. 5). In line with the analysis conducted on the FFPE section alone, the integrated clusters mapped onto defined anatomical structures of the mouse brain including the cerebral cortex, hypothalamus, fiber tracts, thalamus, striatum, lateral ventricle and the hippocampal region. Cluster 11 was more challenging to define by anatomical properties alone but some of the defined marker genes (*Acta2, Tagln, Aqp4* and *Col1a1*) suggested an enrichment of cell types commonly found in the brain vasculature^19^. To exemplify how FFPE-derived expression data can be used for exploratory analyses, we used the integrated clusters as a basis for differential expression (DE) analysis, with the FFPE and FF sections treated independently (Suppl. Fig. 6 and Suppl. Table. 3). The DE analysis demonstrated that a smaller number of marker genes were detected in FFPE section clusters compared to the corresponding FF section clusters (Fig. 1f). However, we found a significant overlap of marker genes between corresponding clusters, where the p-value of the least significant overlap was 1.17*10^−26^ (Suppl. Table. 4). These findings suggest that spatial expression data from FFPE sections can be used to find tissue specific marker genes.

Finally, we used the probabilistic method *stereoscope*^20^ to integrate SMART-seq data, obtained from the Allen Mouse Brain Atlas^21^, with the FFPE dataset. This allowed us to deconvolve the spot transcriptomes into spot-wise proportion estimates of 41 different subclasses present in the SMART-seq data; meaning that the spatial distribution of each cell type could be assessed. Six neuron cell type subclasses were mapped spatially onto layers of the cerebral cortex (Fig. 1g and Suppl. Fig. 7c). The hippocampal region showed highly specific signals for CA1, CA3 and DG cell types, confirming our previous observations from the clustering analysis alone (Suppl. Fig. 7b).Furthermore, we observed how oligodendrocytes were enriched in the fiber tracts (Suppl. Fig. 7d). This has been reported in previous studies and further supports the validity of our results^22^.

To assess the suitability of the method on clinical FFPE samples, we generated spatially barcoded gene expression libraries from FFPE high grade serous ovarian carcinosarcoma (high grade serous carcinoma with a sarcomatous component). Two adjacent sections were prepared respectively from two tissue blocks originating from one biopsy collected from the omentum. H&E stained bright field images revealed a heterogeneous mixture of mainly carcinoma and sarcoma cells interspersed by adipose lobules (Fig 2a) along with immune cell infiltration, vasculature and presence of a few scattered psamomma bodies. The carcinomatous element of the tumor was high-grade serous cancer, presenting a morphology of high cell density nests with numerous irregular slit-like spaces due to fusion of papillae. The sarcomatous element was characterized by fibrous spindle cells with increased nuclear atypia.

We decided to use Non-negative Matrix Factorization (NNMF) to explore our HGSC data in an unbiased manner. In short, this approach models the gene expression as a conical linear combination of a predefined number of factors. In this analysis, we decomposed the gene expression data from these samples into a total of 15 factors which were explored as factor activity maps (Suppl. Fig. 8). The selection of 15 factors was by no means exhaustive, but seemed to recapitulate the major morphological patterns observed in the H&E image. The superposition of the NNMF factor activities onto the histological images showed that factors were indeed spatially distributed and associations to observable histological features could be easily established by a pathologist with expertise on gynecological cancers (Fig. 2a, c). The spatial activity maps were also consistent across consecutive sections, indicating low technical variability (Suppl. Fig. 8).

Factors 1, 7 and 12 were linked to the areas enriched for sarcoma cells by association of high factor activity with histological features of the H&E images (Fig. 2a). Pathway analysis of the top contributing genes for all three factors revealed a strong enrichment for the epithelial to mesenchymal transition (EMT) (Suppl. Fig. 9 and Suppl. Table. 6)^23^, partly defined by the up-regulation of *COL1A1, COL1A2, COL3A1, VIM* and *MMP2* (Suppl. Table. 5 and Suppl. Fig. 8).

Similarly, for factors 2, 6 and 11, we found a strong association with regions enriched for carcinoma, and found KRT7 to be a major contributor consistent with previous pathological assessment of KRT7+ status (Suppl. Table. 5).

Interestingly, factor 4 seemed to be localized to a subgroup of mixed malignant cells, both carcinoma and sarcoma in the vicinity of adipose tissue. Our results indicated that genes associated with factor 4 are myofibroblast associated (e.g. *ACTC1, MYL1, TPM2*) (Fig. 2, Suppl. Fig. 8 and Suppl. Table. 5). Cancer-associated myofibroblasts have been previously described in the literature^24-27^ have been hypothesized to induce other epithelial cells into a proliferative state, a mechanism which normally directs stromal remodeling for regenerative and healing purposes^24^. Myofibroblastic-epithelial interactions have been already described in endometrial cancer^25^.

Factor 3 was found co-localized to the adipose tissue regions and expressed adipose associated pathways (Fig 2c, Suppl. Fig. 9 and Suppl. Table. 6) whereas factor 5 mapped more specifically to the edges of fat lobules resembling mesothelium. Factor 10 showed strong spatial correlation with arterioles and endothelial smooth muscle.

The remaining factors (8, 9, 13) were identified as being enriched for immune-related pathways (Suppl. Table. 6) although with some remarkable differences between the factors. Factor 8 was driven by MHC class II genes, indicating the presence of antigen-presenting cells. In factor 13, we found immunoglobulins *(IGKC, IGHA1)* and *MZB1* among the top contributing genes (Suppl. Fig. 8 and Suppl. Table. 5), suggesting that this factor represents B-cells. Interestingly, factor 13 was detected in areas where damage to adipose tissue and inflammation had occurred (affected adipocytes showing visibly more depleted lipid droplets), indicating where the tumor is infiltrating, destroying and replacing adipose tissue with tumor cells.

In conclusion, we have devised a simple approach which, to our knowledge, demonstrates for the first time genome-wide spatial gene expression profiling in FFPE biospecimens. Data-driven analysis of FFPE tissue from the mouse brain shows that well known anatomical features can be demarcated based on molecular patterns; and spot transcriptomes can be deconvolved into cell type proportions with the integration of single-cell data. These findings encouraged us to explore the applicability of the method within a clinical context where we unraveled the tissue heterogeneity of a carcinosarcoma biospecimen. Despite the sparsity of the data generated from these two tissue types, our results suggest that the performance of the method is sufficient to capture molecular heterogeneity in tissue transcriptomes, opening up the possibility to investigate and explore the rich resource of clinical and research biobanks.

## Acknowledgements

This project was supported by the Swedish Research Council, Swedish Cancer Society, the Swedish Foundation for Strategic Research, Horizon2020 HCA discovAIR, Knut and Alice Wallenberg Foundation (2018.0172), Erling-Persson Family Foundation (HDCA), and Science for Life Laboratory. We would like to thank the National Genomics Infrastructure (NGI) Sweden for providing infrastructure support. We thank 10X Genomics, Annelie Mollbrink, Reza Mirzazadeh, Sami Saarenpää, Ludvig Bergenstråhle and Mengxiao He for advice and helpful discussions.

## Author contributions

EGV led the experimental work and EGV and LK performed the experiments. LL analyzed the data and LL and EGV prepared the figures. AA developed the single cell mapping method and helped with data analysis. JC designed the clinical experiments. JL supervised the research. EGV, LL and JL wrote the manuscript with input from all authors.

## Ethics declarations

Competing Interests: JL, EVG, LL, LK, AA are scientific advisors to 10x Genomics Inc, which holds IP rights to the ST technology.

## Methods

### FFPE Mouse Brain sample

Male, 25 g, 8-12 weeks of age. Sample number C57BL6J (Adlego Biomedical). Extracted under ethical permit number 4570-2019. RIN 2.9, DV200 65%.

### FFPE Gynecological Carcinosarcoma

Female. High grade serous ovarian carcinosarcoma metastasis to the omentum. Extracted under ethical permit number 2018/2264-31.Untreated. Two regions: 1919-1 RIN 2.5, DV200 69% and 1919-2 RIN 2.30, DV200 64%.

### Sectioning, deparaffinization, and staining of FFPE samples

Gynecological carcinosarcoma microtome sections (12 µm thick) were placed on Visium Spatial Gene Expression slides (10x Genomics) after floating on a water bath at 43 °C. The same was done for the FFPE coronal Mouse Brain section (10 µm thick). After sectioning, the slides were dried at 40°C in a hybridization oven (HB-1000 Hybridizer, LabRepCo) for 1 hour and 45 min. The slides were then placed inside a slide mailer, sealed with parafilm, and left overnight in a refrigerator at 4°C. The slides were deparaffinized by immersion in the following reagents: Xylene (#28975.291 VWR) 7 min twice, EtOH 99% (#84835.290 VWR) 2 min twice, EtOH 96% (#20823.290 VWR) 2 min twice. Staining was performed according to the Methanol Fixation, H&E staining & Imaging for Visium Demonstrated Protocol CG000160 RevA, 10x Genomics^1^. Step 1.3 of the protocol was performed with the following modifications: Step 1.3.q slides were incubated in Dako bluing buffer (#CS70230-2 Agilent) for 30 s. After Step 1.3.z, slides were mounted with 200 µl glycerol 85% and a cover glass was applied on the slide.

### HE imaging

Slides were scanned under a high-resolution microscope Metafer Slide Scanning Platform (Metasystems) to obtain tissue tile images and software VSlide (Metasystems) to stitch the high-resolution images together. After imaging, the glycerol and cover glass were carefully removed by holding the slides in an 800 ml water beaker and letting the glycerol diffuse until the cover glass detached and density changes were no longer visible in the water. The slides were then dried at 37°C.

### Analyte retrieval and permeabilization

Slides were mounted in an ArrayIt metallic hybridization cassette (#AHC1×16 ArrayIt). A collagenase mix (986 µl HBSS buffer (#14025-050 Life Technologies), 10 µl BSA (#B9000S Bionordika), 4 µl collagenase I (#17018-029 Life Technologies)) was equilibrated to 37°C and then 75 µl were added to each of the wells in the cassette. The slides were sealed and incubated for 20 min at 37 °C in a Thermoblock (ThermoMixer with Thermoblock, Eppendorf) with heated lid (ThermoTop, Eppendorf). Once the incubation was complete, the collagenase mix was pipetted off and the slides were washed by with 100 µl of 0.1 x SSC buffer (#S6639-1L SigmaAldrich, diluted in RNAse DNAse free MQ) in each well. Subsequently, 100 µl of TE buffer pH 8.0 (#AM9849 ThermoFisher) was added, and the slides were sealed and incubated for 1 hour at 70°C in a Thermoblock with heated lid. After the incubation, the slides were taken out of the Thermoblock and left to equilibrate at room temperature for 5 min. Meanwhile, 0.1% pepsin solution (P7000-25G SigmaAldrich) dissolved in 0.1M HCl (#318965-1000ML SigmaAldrich) was equilibrated to 37°C. After the incubation and one wash per well with 100 µl 0.1 x SSC buffer, permeabilization of the tissue was carried out by adding 75 µl pepsin solution, sealing the slides and incubating with heated lid at 37°C for 30 min. The following steps were performed according to the Visium Spatial Gene Expression User Guide CG000239 Rev C, 10x Genomics^2^. Reverse Transcription was performed as described in Step 1.2 of the User Guide, with the difference that the slides were still masked in ArrayIt metallic hybridization cassettes instead of Visium Slide cassettes.

All step names and numbers cited in the document correspond to the Visium Spatial Gene Expression User Guide CG000239 Rev C unless specified otherwise.

### Second Strand synthesis and denaturation

Second strand synthesis and denaturation were carried out as described in the standard Visium Spatial Gene Expression User Guide Step 2 with the difference that the slides were masked in ArrayIt metallic incubation chambers instead of Visium Slide cassettes.

### cDNA amplification, cleanup and quantification

cDNA derived from mouse brain and carcinosarcoma tissue was amplified by 14 cycles of PCR with settings specified in Visium Spatial Gene Expression User Guide Step 3.2. Sample cleanup was performed using 0.8x SPRIselect beads instead of 0.6x, meaning that 80 µl of SPRIselect beads were added to 100 µl sample instead of 60µl on Step 3.3.a. The Buffer EB volume for eluting was reduced to 15 µl instead of 40.5 µl in Step 3.3.j of the protocol. The cDNA yield was checked by using 1µl sample for quality control on a BioAnalyzer High Sensitivity chip (Agilent) as in Step 3.4.

### Fragmentation, end repair and A-tailing

A total of 10 µl of each sample were fragmented in a thermal cycler as described in step 4.1 with the alteration that fragmentation run time was reduced to 1 min instead of 5 min. Samples were cleaned up post-fragmentation using 0.8x SPRIselect beads. 40 µl SPRIselect beads were added to 50 µl sample and incubated at room temperature for 5 min. The tubes were then placed on the high magnet position until the solution cleared. Supernatant was discarded and the beads were washed with 125 µl 80% ethanol for 30 s two times. After removing the 80% ethanol, beads were allowed to dry on the magnet for 2 min (avoiding over-drying) followed by addition of 50.5 µl of Buffer EB to each sample.

The samples were removed from the magnet, vortexed, and centrifuged briefly to mix the beads with Buffer EB and were then incubated for 2 min at room temperature. After incubation the samples were placed on the low magnet position until the solution cleared. 50 µl of each sample was transferred to a new tube strip.

### Adapter ligation SPRIselect post-ligation clean-up

These steps were performed as described in Visium Spatial Gene Expression Reagent Kits User Guide CG000239 Rev C (10x Genomics)^2^, Steps 4.3 and 4.4.

### Sample index PCR and clean-up

Samples were indexed as in Visium Spatial Gene Expression Reagent Kits User Guide, Step 4.5, with 8 cycles of PCR. Subsequently, samples were cleaned up as in Step 4.6 with the difference that in step 4.6.m, 40.5 µl Buffer EB was added for elution instead of 35.5 µl, thus 40µl being the final eluted volume that was transferred to a new tube strip. An additional 0.8x SPRIselect bead clean-up step was performed. 32 µl SPRIselect beads were added to 40 µl sample and incubated at room temperature for 5 min. Then the tubes were placed on the high magnet position until the solution cleared. Supernatant was discarded and the beads were washed with 200 µl of 80% ethanol for 30 s twice. After removing the 80% ethanol, beads were allowed to dry on the magnet for about 2 min (avoiding over-drying) followed by addition of 15 µl of Buffer EB to each sample. The samples were removed from the magnet, vortexed, and centrifuge briefly so the beads were mixed with Buffer EB, then incubated for 2 min at room temperature. After incubation the samples were placed on the low magnet position until the solution cleared. Eluted samples were transferred to a new tube strip.

### Post-library QC and dilution

Performed as in Step 4.7 in the Visium Spatial Gene Expression Reagent Kits User Guide. In addition, 2 µl of each sample were used for dsDNA HS Qubit assay (Thermo Fisher Scientific) to determine sample concentration.

### Sequencing

Libraries were sequenced using Illumina’s Nextseq 500. A total of 4 samples per run, with 75 cycles High Output kits, with pair-end, dual index sequencing and custom primer for Read 2. (Integrated DNA Technologies, Sequence AAG CAG TGG TAT CAA CGC AGA GTA CAT GGG, Purification HPLC). A total of 2 ml of 0.3 µM custom primer for Read 2 were loaded into well number 8 of the sequencing cartridge. Libraries loading concentration was 1.8 pM with a 1% PhiX spike-in. Read 1: 28 cycles, i7 index: 10 cycles, i5 index: 10 cycles, Read 2: 44 cycles.

### Data pre-processing

The raw fastq files containing the cDNA sequences (R2) were pre-processed to remove TSO primer sequences and poly(A) homopolymers using *cutadapt* (v2.8)^3^. TSO sequences were trimmed by defining the TSO sequence as a non-internal 5’ adapter (which will remove partial or full TSO sequences from the 5’ end) with a minimum overlap of 5 bp and an error tolerance of 0.1. Poly(A) homopolymers were trimmed by defining 10 A’s as a regular 3’ adapter (which will remove stretches of poly(A) found anywhere in the sequence as well as the trailing base pairs with a minimum overlap of 5bp). To search for and trim both adapter types from the same read sequences we set the --times option to 2.

### Data processing

All paired fastq files (after TSO and poly(A) trimming) were processed using spaceranger v1.0.0 together with the corresponding Hematoxylin and Eosin (H&E) stained images in jpeg format. For mapping of the data, we used the mm10-3.0.0 *mus musculus* reference genome for mouse samples and the GRCh38-3.0.0 *homo sapiens* reference genome for human samples (both included in the Space Ranger distribution v1.0.0).

### Sample selection

With the developed FFPE protocol we generated four mouse brain libraries, of these one generated sufficient sequencing information. This library was used for the gene expression analysis and single cell mapping effort, described in this study. With the FFPE protocol applied on clinical samples, using a set of four patient samples, we generated sufficient data from two samples (in duplicates). The well recognized variation between archival blocks in terms of RNA accessibility and stability will require new investigations to identify relevant QC metrics for spatial barcoding schemes that builds on the use and the accessibility of the poly (A) tail of mRNA instead of rRNA integrity number. The integrity and accessibility of the poly (A) signature is currently not addressed in the standard operating procedures for FFPE samples (http://www.isber.org; http://biospecimens.cancer.gov).

### Analysis of mouse brain data

The Fresh Frozen (FF) Visium gene expression dataset (coronal section of one hemisphere of the mouse brain) was downloaded from 10x Genomics website^4^. Using the Allen Brain Atlas as reference, the dataset FF section was determined to have been collected approximately –2.2 mm from Bregma along the anterior-posterior axis. The FFPE section (coronal section of one hemisphere) was collected at approximately the same coordinate from Bregma along the anterior-posterior axis. Morphology of the relative location from Bregma was checked under the microscope every ∼100 µm apart sections.

### Registration of H&E images to the Allen Mouse Brain Atlas anatomical reference

The H&E images of the FFPE and FF tissue sections were registered to the Allen Brain Atlas anatomical reference using the *wholebrain* (v0.1.1) framework in R. To summarize, the H&E images were first converted into grayscale and the intensity values inverted to give the Hematoxylin stained nuclei higher intensity values than the background. These inverted H&E images were then used as input for cell segmentation and registration to the anatomical reference (−2.2 mm from Bregma) by manual addition of correspondence points, i.e. points matching anatomical features of the H&E image and the anatomical reference.

### Analysis of FFPE mouse brain tissue

The FFPE Visium data was filtered by (1) removal of spots with fewer than 100 unique genes, (2) removal of mitochondrial protein coding genes and (3) removal of genes annotated to non-coding RNA biotypes (“antisense”, “lincRNA”, “pseudogenes”). All three filtering steps were computed within an R programming environment using the *STUtility* package (v1.0)^5^ and the filtered expression matrix was converted to a Seurat object. Normalization, dimensionality reduction, clustering, UMAP embedding and DE analysis was conducted using the *Seurat* R package; normalization – SCTransform() with variable.features.rv.th = 1.1, variable.features.n = NULL and return.only.var.genes = FALSE; dimensionality reduction – RunPCA(); UMAP embedding – RunUMAP() with dims = 1:25, n.epochs = 1,000 and n.neighbors = 30; clustering – FindNeighbors() with dims = 1:25 and FindClusters() with the resolution set to 1.4; DE analysis - FindAllMarkers(). The DE analysis was computed using a pairwise Wilcoxon Rank Sum test between the spots in each cluster contrasted to all other spots in the dataset. Only genes with a positive avg_logFC value and an adjusted p-value lower than 0.01 were kept in the analysis.

### Visualization of marker gene expression by ISH

One candidate marker gene was selected from each cluster defined in the FFPE dataset and queried in the ISH data explorer of the Allen Mouse Brain Atlas. The selection of marker genes was done manually based on adjusted p-value, high avg_logFC values and also based on presence in the ISH atlas. ISH data (microscopy images) of coronal tissue sections were selected from the data explorer at a distance from Bregma along the anterior-posterior axis matching approximately that of the FFPE and FF tissue sections (−2.2mm) and downloaded in full resolution from the High-Resolution Image Viewer. Gene expression patterns (normalized expression) from the FFPE Visium data were visualized side by side with the ISH images.

### Integrated analysis of FFPE and FF mouse brain tissue sections

The FFPE and FF coronal section datasets were first merged by gene ID followed by filtering, normalization with SCTransform() and dimensionality reduction with RunPCA() as described in the previous section for the FFPE dataset. Biotype annotations were extracted from the GENCODE reference GTF file (GRCh38 genome assembly), shipped with the spaceranger command line tool. These biotype annotations were then used to group genes into biotype groups. In addition, mitochondrial genes (prefixed mt-in MGI nomenclature) were defined as “protein coding mitochondrial” and ribosomal protein genes (prefixed Rpl or Rps in MGI nomenclature) were defined as “protein coding ribosomal”. Within each condition (FFPE and FF) the relative amounts of molecules found within each biotype were computed by aggregating all UMI counts within each biotype group and dividing by the total number of UMI counts.

Gene attributes were computed within each condition (FFPE and FF) by summation of UMI counts for each gene across all spots (bulk level) followed by log10-transformation (pseudocount 1). The log-transformed values were then used as input for the calculation of a Pearson correlation score using the stat_cor() function from the *ggpubr* R package.

Next, we applied the harmony integration method on the two datasets (FFPE and FF) to generate a dimensionality reduction representation (embedding) of spot transcriptomic profiles with the RunHarmony() function from the *harmony* R package. This embedding was then used as input for clustering – FindNeighbors() with dims = 1:30 and FindClusters() with the resolution set to 0.7 and UMAP – RunUMAP() with dims = 1:30 and n.epochs = 1,000. After integration and clustering, the dataset was split by condition into two new subsets, i.e. FFPE and FF. A DE test was computed using FindAllMarkers() on the integrated clusters in each subset. DE genes with a positive avg_logFC value and an adjusted p-value lower than 0.01 were kept, which defined the set of markers for each integrated cluster in each condition (FFPE or FF). To estimate the significance of the overlap of marker genes across conditions, we used the testGeneOverlap() function from the *GeneOverlap* R package^6^. For each cluster, assume that the gene sets A and B represent the marker genes detected in FFPE and FF. The problem is formulated to test whether the variables A and B are independent by constructing a contingency table followed by Fisher’s exact test.

### Cell type mapping with stereoscope

To assess how certain cell types were distributed in the tissue sections (both FF and FFPE) we used the tool *stereoscope* (v.3.0)^7^ to integrate scRNA-seq data obtained with the SMART-seq protocol^8^. The area of a capture location (spot) in the spatial assay is large enough to host several cells; a common estimate is 1-10 cells per spot. Hence, the observed gene expression at each spot can be considered a mixture of contributions from multiple cells. Furthermore, the cell population associated with a spot is not necessarily homogeneous, meaning different cell types may be represented and a one-to-one relationship between spot and cell type is not guaranteed. Thus, to make an informed statement regarding certain cell types’ arrangement within the tissue, based on the spatial transcriptomics data, one must first *deconvolve* the gene expression profiles. The method we used, *stereoscope*, addresses the issue of mixed contributions by leveraging “pure” single cell data (one datapoint represents one cell and type) to first characterize the expression profile of each cell type and then estimate the composition of cell types that best explain the observed gene expression at each spot using these profiles. It is a probabilistic approach, where both single cell and spatial data are modelled as negative binomial distributed. The output from *stereoscope* is a matrix in the format of [n_spots]x[n_types] where each element represents a proportion estimate (not a score) of a given cell type at a specific spot. Any cell type with less than 25 members was excluded from the analysis, all cells from types with more than 25 but less than 500 members were used, 500 cells were randomly sampled from types with more than 500 members. A batch size of 2,048 and 75,000 epochs were used in both steps of the procedure. We used the 5,000 most variable genes (in the single cell data) in the analysis.

### Filtering and normalization of carcinosarcoma datasets

Raw expression matrices from the four gynecological carcinosarcoma tissue samples were first merged and then filtered to (1) remove spots with fewer than 150 unique genes, (2) remove genes with fewer than 10 counts across the whole dataset and (3) remove mitochondrial protein coding genes as well as all non-coding RNA biotypes (“antisense”, “lincRNA”, “pseudogenes”). All three filtering steps were computed within an R programming environment using the *STUtility* package and the filtered expression matrix was converted to a Seurat object. The raw counts were normalized using the Variance Stabilizing Transformation (VST) method implemented in the SCTransform function in the *Seurat* package (v3.1.5) (settings: return.only.var.genes = FALSE, variable.features.n = NULL, variable.features.rv.th = 1.1).

### Non-negative Matrix Factorization and pathway analysis

The VST normalized and scaled expression matrix was decomposed into 15 factors using a Non-negative Matrix Factorization method from the *NNLM* package modified as described by Wu et al.^9^ and implemented in the RunNMF function in *STUtility*. In summary, the normalized and scaled gene expression matrix (A) is first transformed to contain strictly non-negative values and is then decomposed into two matrices W*H, where W is the (genes x samples) gene loadings matrix and H is the (factors x samples) spot embeddings matrix. From the gene loadings matrix, we selected a set of top contributing genes for each factor to use as input for pathway analysis using a simple mean standard deviation method. Each factor gene loading vector was first log-transformed to produce a new vector x, and genes with a value higher mean(x) + 1.645*sd(x) were kept as a set of “top contributors” for the factor. Each set of “top contributor” genes were used as input for pathway analysis using the gost() function from the *gprofiler2* R package with two different sources; *GO:BP* and the “cancer hallmark collection” available from MSigDB^10^.

## Supplementary Figures

**Supplementary fig. 1.**
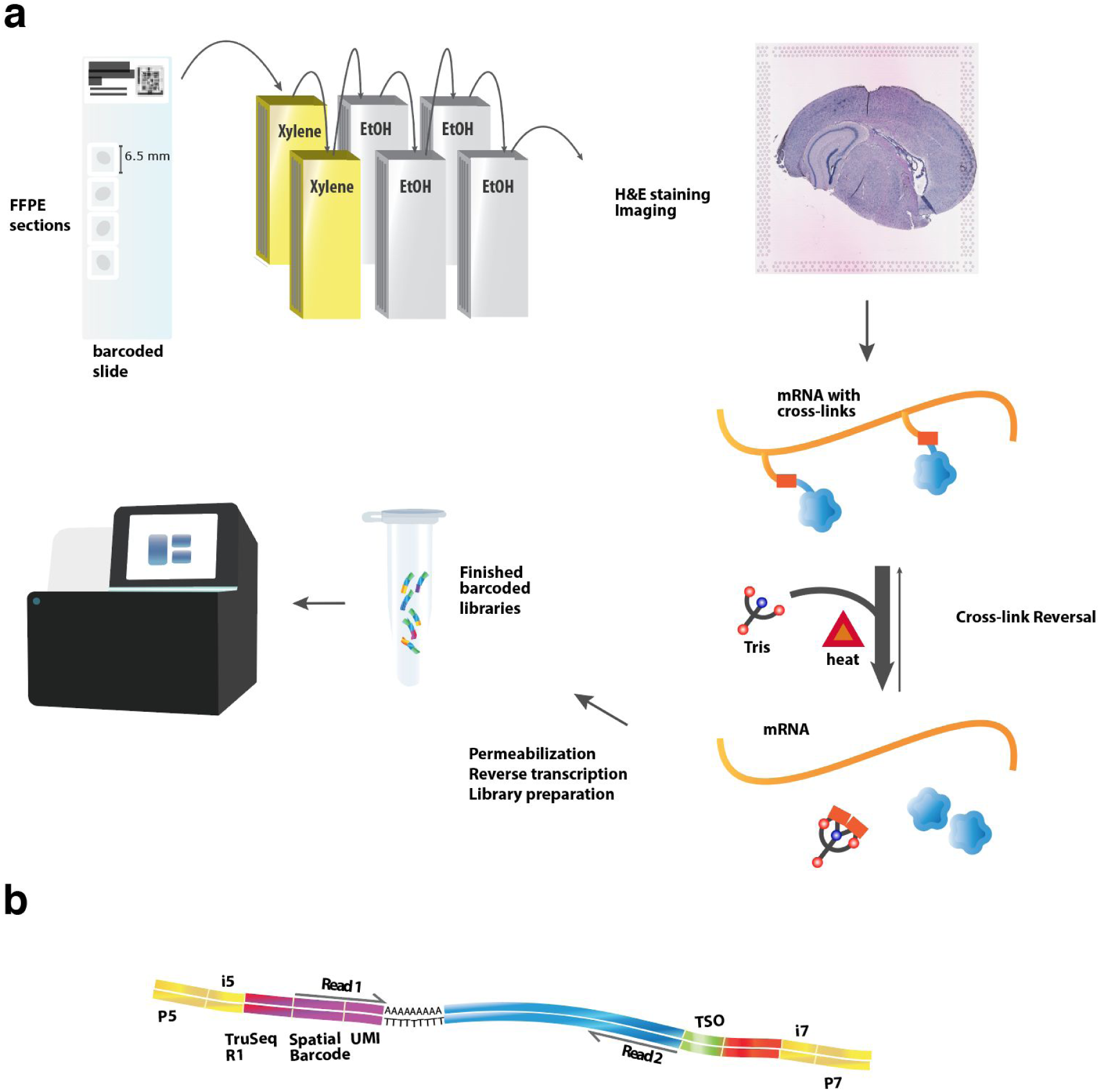
Visualization of mRNA expression across FFPE tissue sections with adapted protocol. **a**, Tissue sections are placed on spatial gene expression slides. The slides are deparaffinized following a standard protocol (see methods). H&E stained sections are imaged under a high-resolution microscope. Cross-links are reversed by heat and in the presence of Tris-EDTA buffer at pH 8.0. Tissues are permeabilized so that mRNA diffuses to the barcoded surface probes and hybridizes with its oligo(dT) capture region. A cDNA strand is synthesized by reverse transcription. Finished libraries are sequenced with custom primer for read 2 reverse complementary to template switch oligo (TSO) sequence in the final construct. **b**, Final construct overview.

**Supplementary fig. 2.**
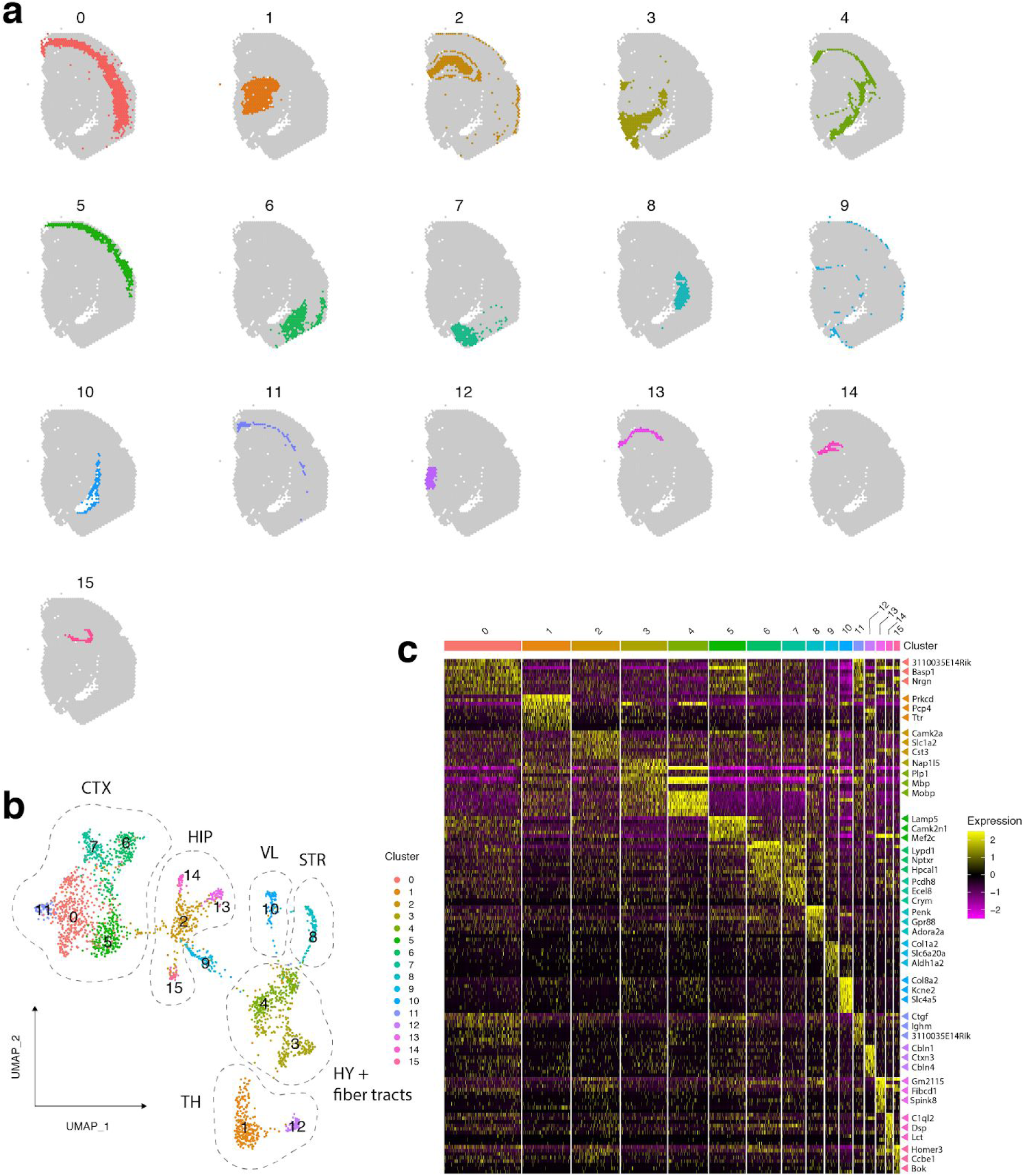
Clustering and marker detection in a coronal section from FFPE mouse brain tissue. **a**, Spatial mapping of 16 clusters on tissue section spot coordinates. **b**, 2D UMAP embedding of spatial gene expression data colored by cluster. Clusters 2, 13, 14 and 15 form the hippocampal region (HIP) where cluster 13 maps to field 1 of the pyramidal layer (CA1sp), cluster 14 maps to the dentate gyrus (DG-sg) region and cluster 15 maps to field 3 of the pyramidal layer (CA3sp) region. Cluster 2 shows a less specific pattern in **a** where it makes up the majority of the HIP and also part of the outermost layer of the cerebral cortex (CTX). This can also be observed in the UMAP where cluster 2 connects the HIP to CTX. Clusters 0, 5, 6, 7 and 11 form the CTX which can be further divided into cortical subplate and olfactory areas (6, 7) and the isocortex (0, 5, 11). Other examples include: hypothalamus (HY) – cluster 3; fiber tracts – cluster 4; thalamus (TH) – clusters 1 and 12; striatum (STR) – cluster 8; lateral ventricle (VL) – cluster 10. Cluster 9 maps mostly to the edges of the CTX and is enriched for genes expressed primarily in vascular tissue. **d**, Heatmap of differentially expressed genes within each cluster, represented by normalized and scaled expression values. Three selected genes per cluster with a high significance (adjusted p-value < 0.01) are highlighted for each cluster on the right side of the heatmap.

**Supplementary fig. 3.**
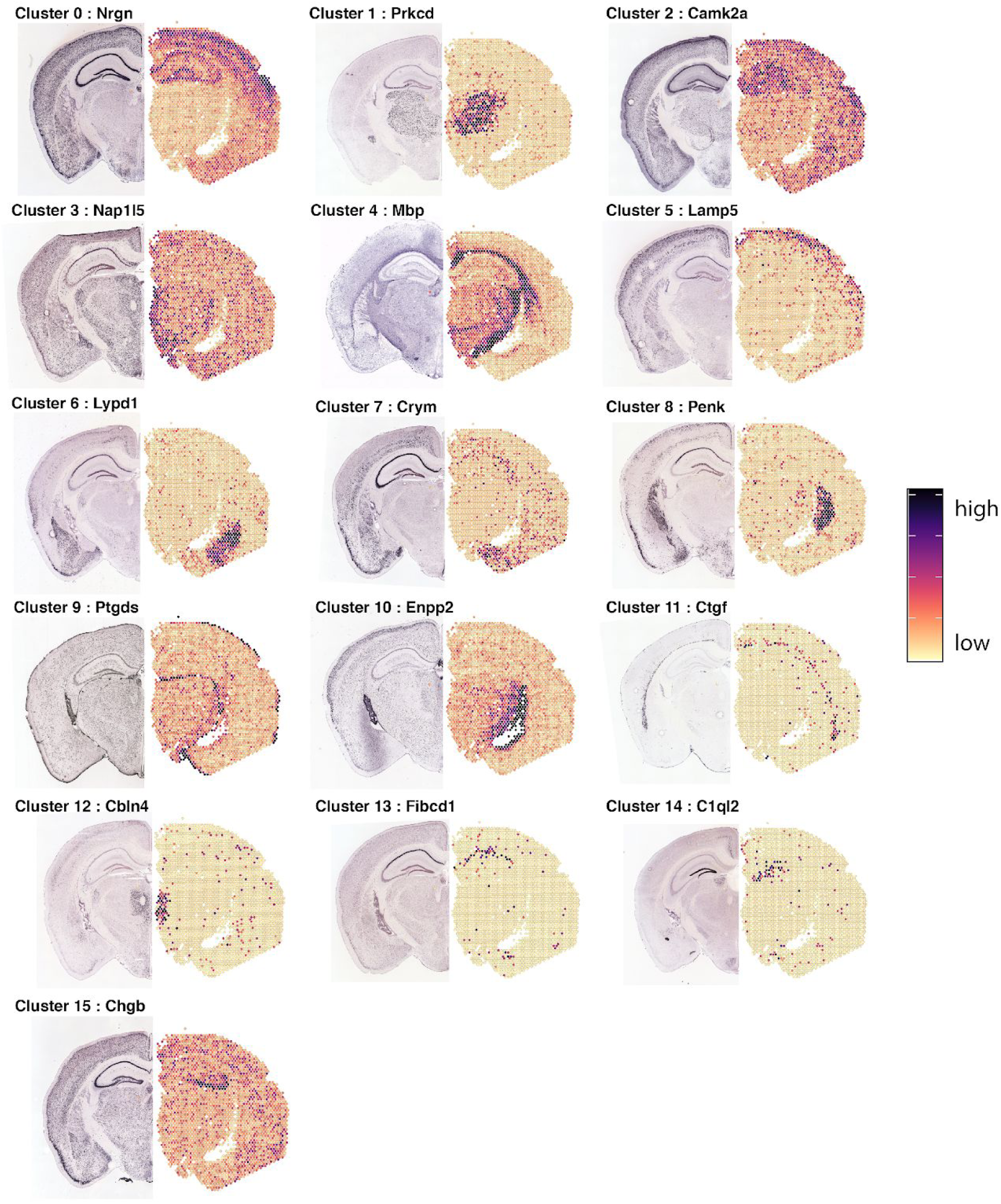
Validation of marker genes by ISH. Selected marker genes visualized by *in situ* hybridization (ISH) from the Allen Mouse Brain Atlas (left hemisphere) versus scaled expression values from our FFPE data (right hemisphere). Marker genes were selected based on significance (adjusted p-values) and average log-fold change from the differential expression analysis.

**Supplementary fig. 4.**
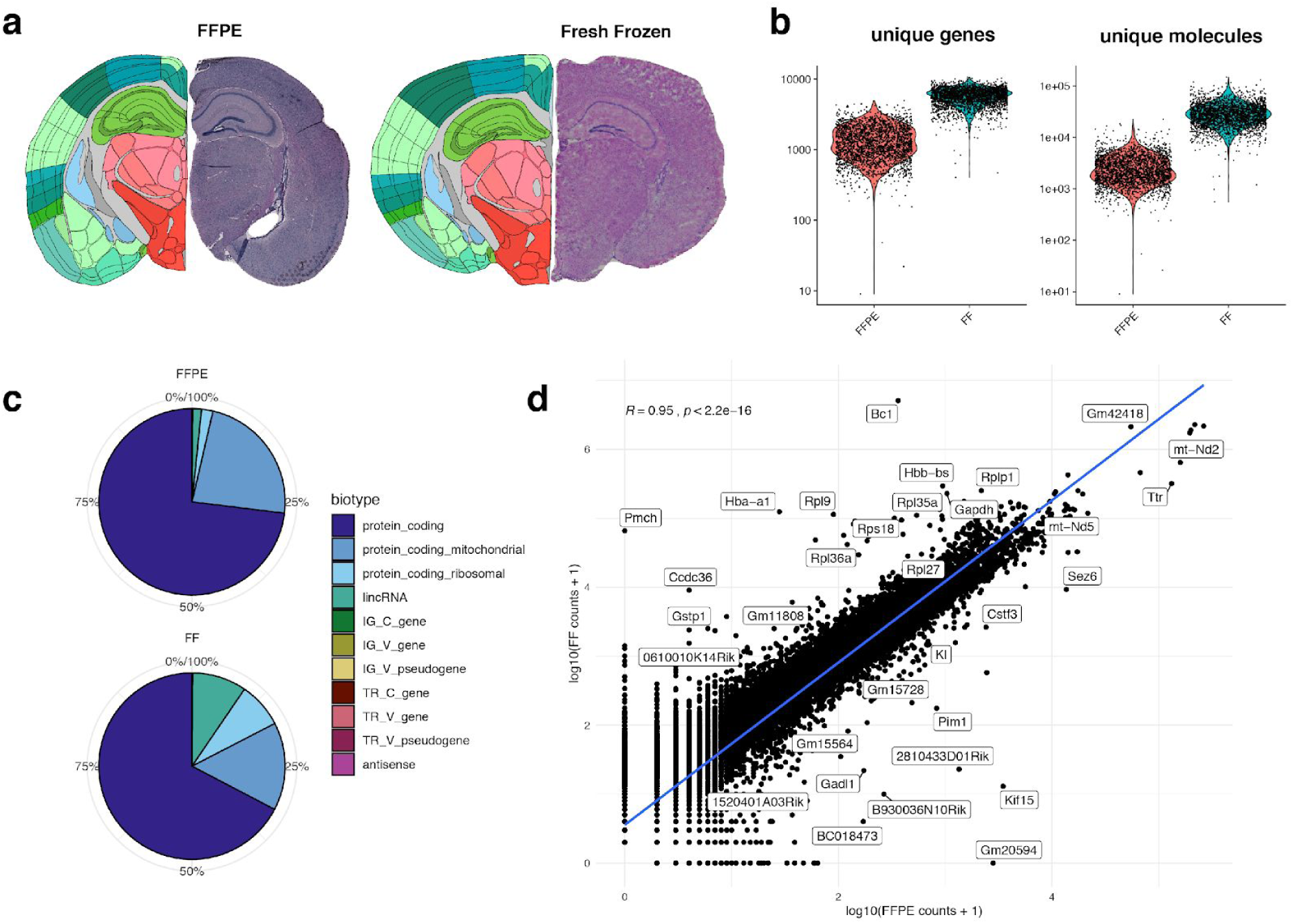
Image registration, quality control and correlation of FFPE and FF tissue sections. **a**, H&E images of the FFPE and Fresh Frozen (FF) tissue sections (right hemisphere) next to the registered anatomical reference from the Allen Mouse Brain Atlas (left hemisphere). The registration was done using the *wholebrain* framework with the stereotactic coordinate set to -2.2 from bregma along the anterior-posterior axis for both tissue sections. **b**, Quality metrics shown as violin plots for the FFPE and FF sections (y-axis in log-scale). The median number of unique genes per spot was ∼1200 and ∼6000 respectively. The median number of unique molecules (UMIs) per spot was ∼2200 for the FFPE section and ∼27200 for the FF section. **c**, Composition of RNA biotypes in the two tissue sections. Notably, the FFPE section contained a higher fraction of mitochondrial protein coding genes and a lower fraction of both ribosomal protein coding genes as well as lncRNA. **d**, Scatter plot showing the correlation between log10-transformed UMI counts per gene for two tissue sections with a Pearson correlation coefficient of 0.95. Highlighted outlier genes indicate that hemoglobin genes *Hba-a1* and *Hbb-bs* and certain ribosomal protein genes are more highly expressed in the fresh frozen tissue section.

**Supplementary fig. 5.**
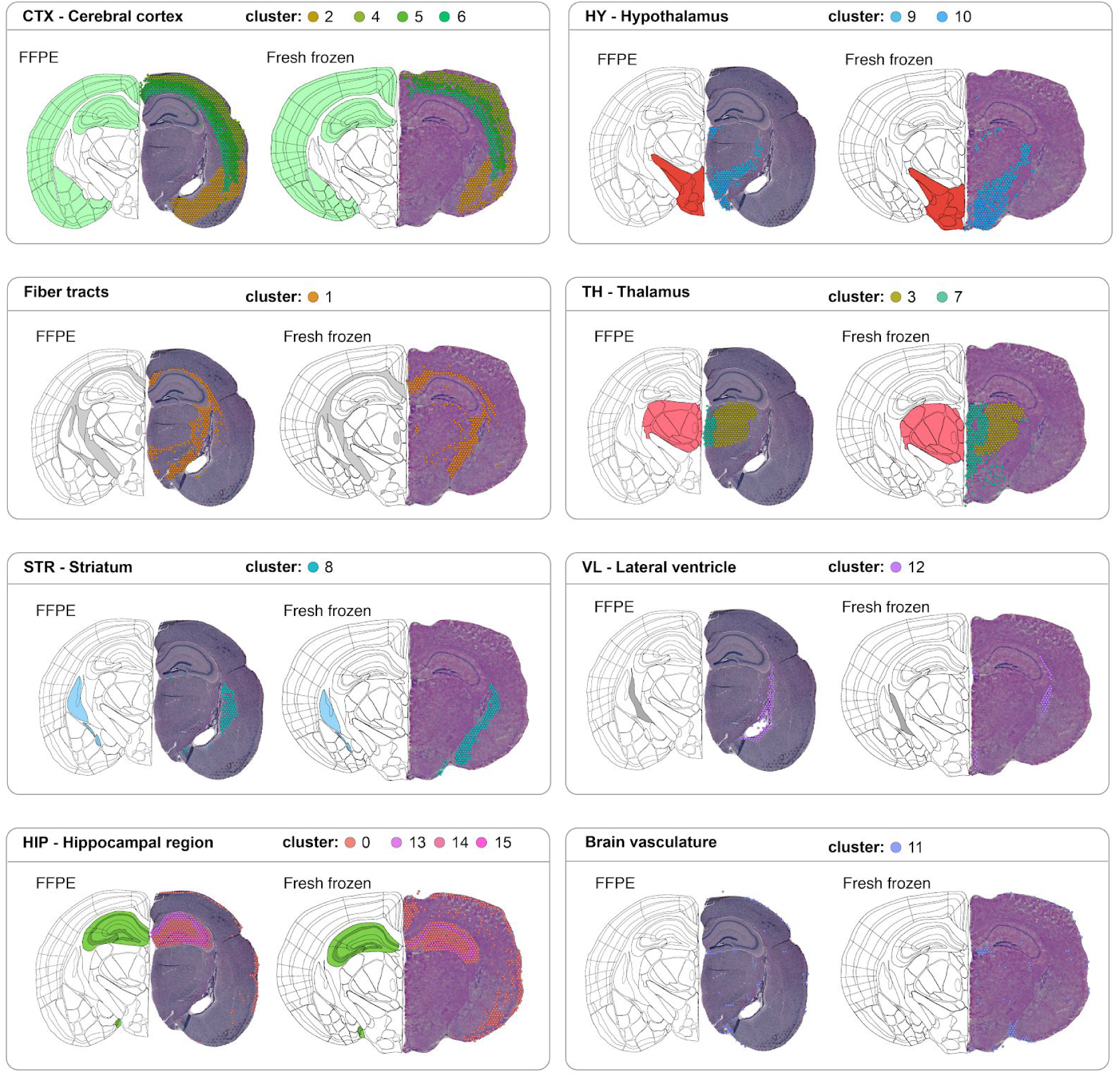
Integrated clusters across FFPE and FF tissue sections. The FFPE and Fresh Frozen datasets were integrated and clustered using *harmony*. Clusters have been grouped together based on their co-localization with anatomical landmarks from the Allen Mouse Brain Atlas.

**Supplementary fig. 6.**
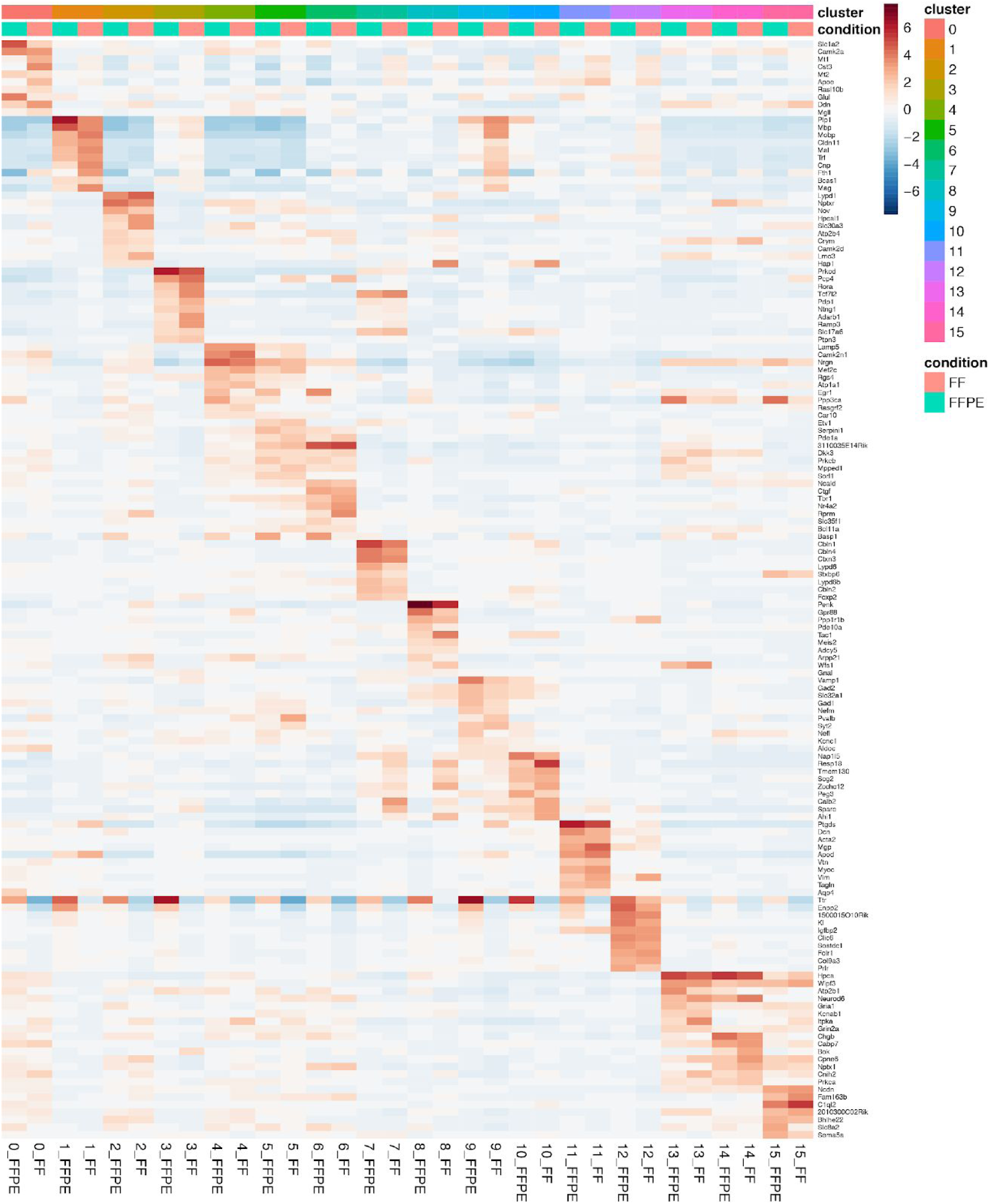
Integrated cluster marker genes detected in both FFPE and Fresh Frozen data. Heatmap showing the averaged scaled expression of marker genes detected in each integrated cluster which are found in both tissue sections. Markers were extracted using a DE analysis test applied independently on the FFPE and Fresh Frozen (FF) datasets where the spots within each cluster were contrasted to all other spots. Each cluster is displayed in two columns; FFPE and FF. For clusters with more than 10 shared marker genes, only 10 are shown.

**Supplementary fig. 7.**
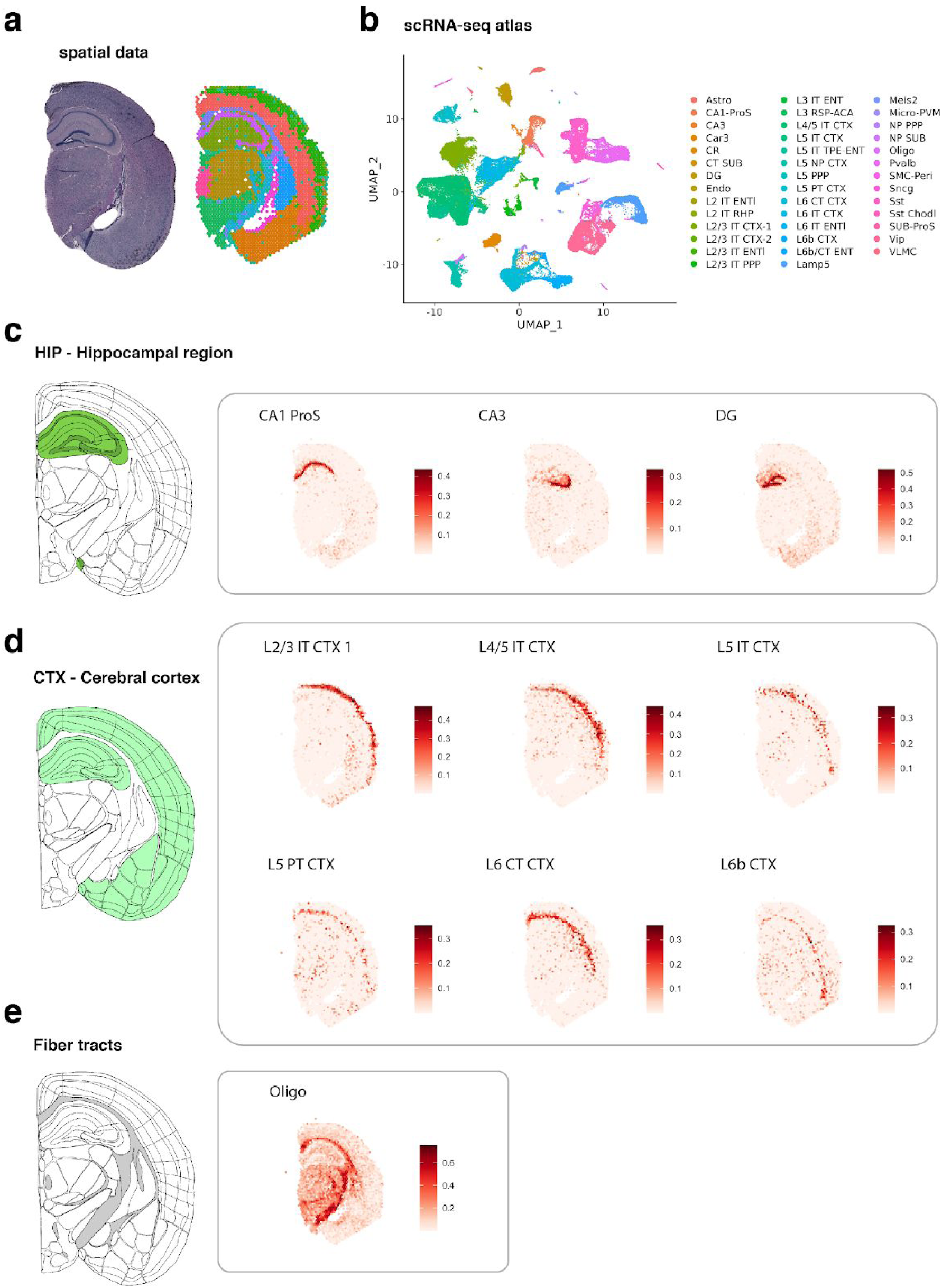
Cell type subclass mapping with stereoscope. **a**, Clusters defined in spatial gene expression data obtained from FFPE coronal tissue section **b**, UMAP visualization of the SMART-seq scRNA-seq data from the Allen Brain Atlas used for cell type mapping with *stereoscope*, with a total of 41 subclasses. **c, d, e**, A selected subset of cell type proportions co-localized with the hippocampal region (HIP), cerebral cortex (CTX) and fiber tracts. Color bars represent cell type proportions. **c**, Three distinct cell types co-localized with the HIP; CA1 ProS mapped to field 1 of the pyramidal layer (CA1-sp); CA3 mapped to field 3 of the pyramidal layer (CA3-sp) and DG mapped to the dentate gyrus (DG-sg). **d**, Six cell types mapped to layers of the CTX. **e**, Oligodendrocytes mapped to the fiber tracts.

**Supplementary fig. 8.**
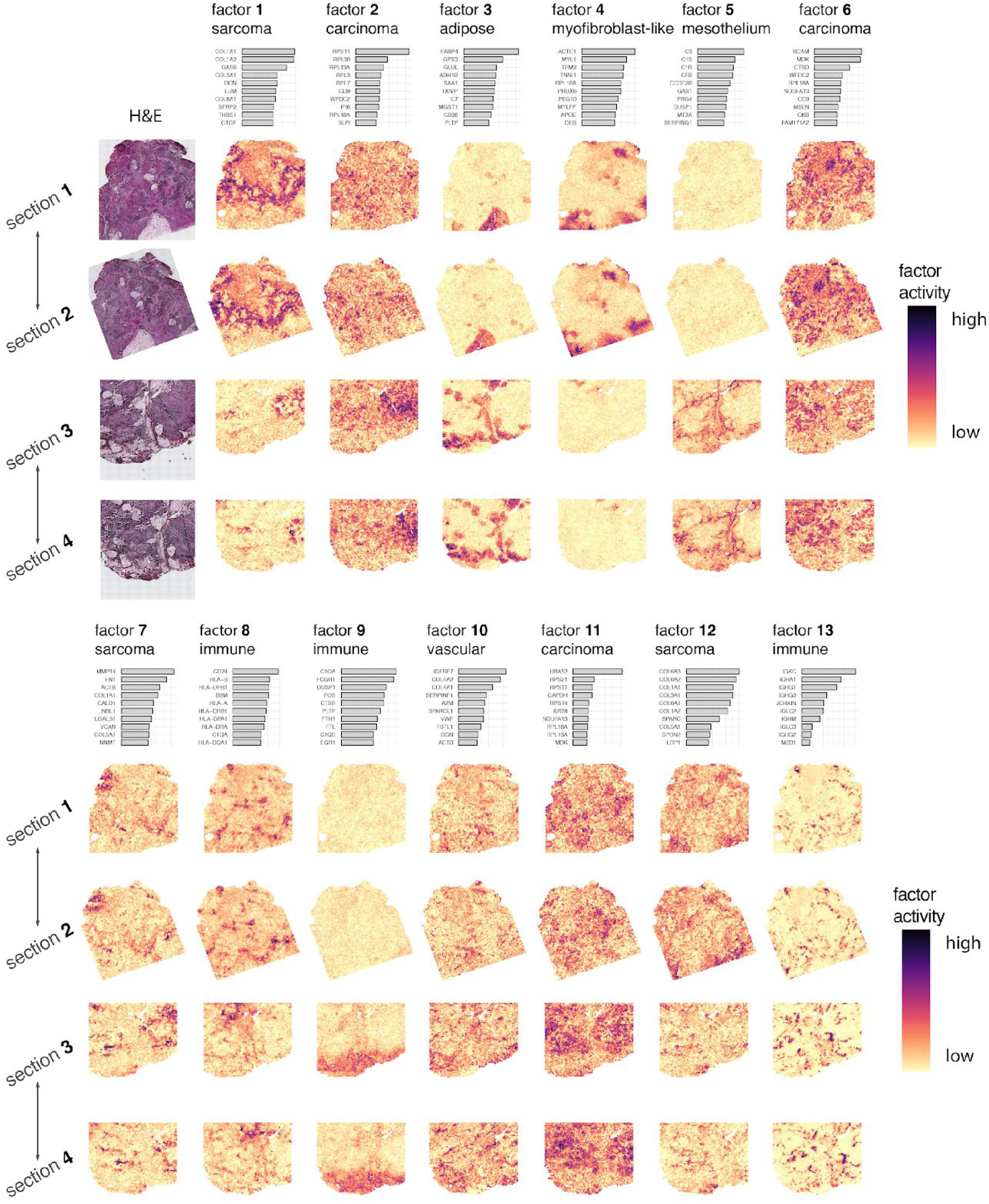
Spatial factor activity maps of four carcinosarcoma tissue sections. The first column shows the H&E images linked to each spatial activity map. The remaining columns represent factor activity maps and consist of four rows, with one row for each tissue section. Adjacent sections are indicated by double headed arrows. The bar charts on top of each activity map shows the top 10 contributing genes for each factor. Factors 14 and 15 were driven mainly by ribosomal protein coding genes and were therefore omitted.

**Supplementary fig. 9.**
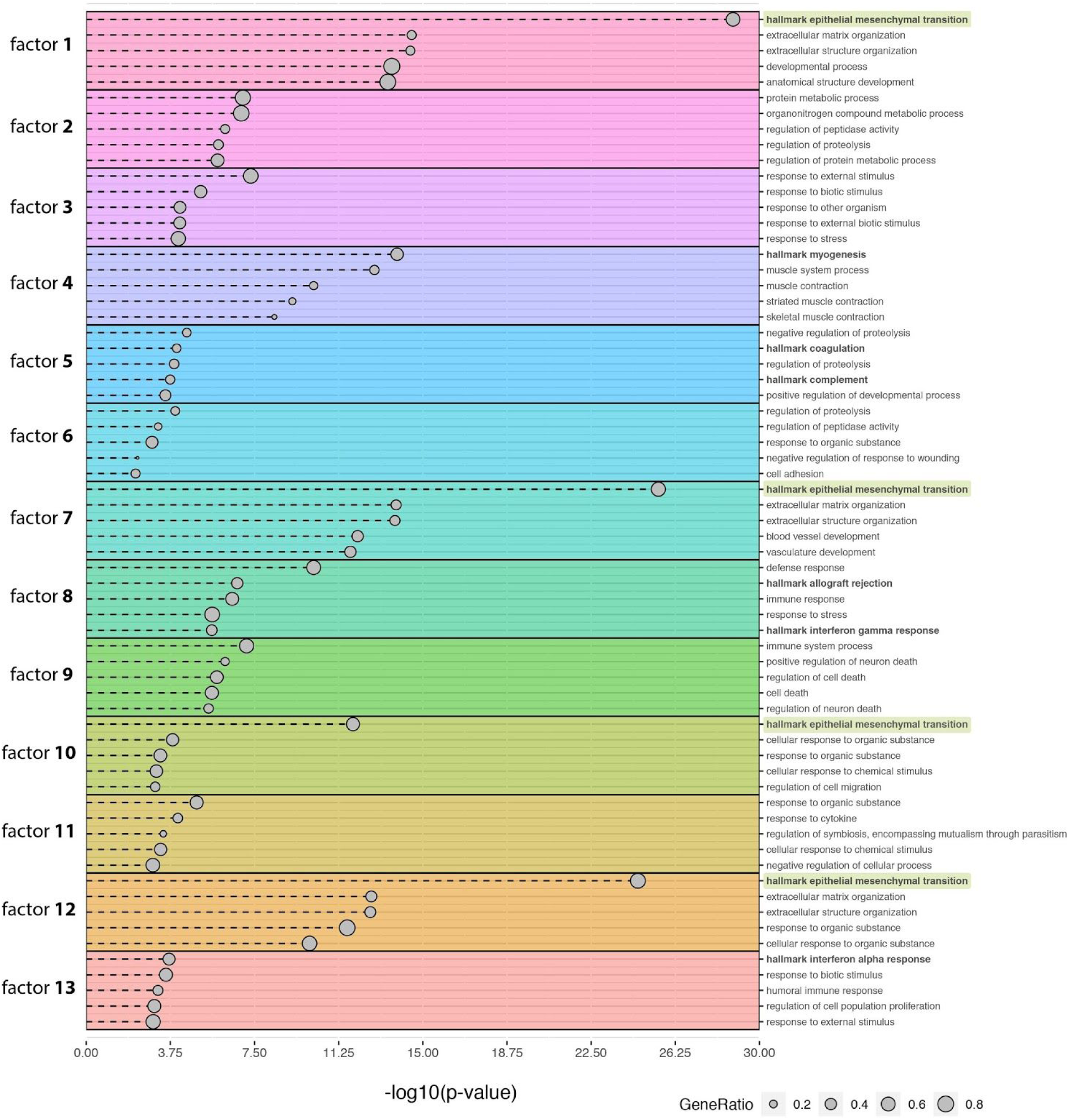
Pathway analysis of factors in carcinosarcoma tissue. Top 5 most significant pathways per factor based on gene ontology (biological processes) and the cancer hallmark gene set collection from MsigDB. Terms from the hallmark gene set collection are highlighted with bold text with the epithelial to mesenchymal transition (EMT) term highlighted with green boxes. The GeneRatio is defined as the number of genes shared between the top contributing genes per factor and the term gene set divided by the total number of top contributing genes.

